# Anabolic lipid metabolism regulates adipose type 2 innate lymphoid cell differentiation to maintain metabolic health

**DOI:** 10.1101/2024.03.26.586766

**Authors:** Maria Rafailia Theodorou, Jiangyan Yu, Fabian Nikolka, Jelena Zurkovic, Chantal Wientjens, Patricia Weiss, Roman Rombo, Fotios Karagiannis, Christoph Thiele, Jan Hasenauer, Karsten Hiller, Christoph Wilhelm

## Abstract

Group 2 innate lymphoid cells (ILC2) residing in the adipose tissue play an important role in maintaining the metabolic health and energy balance of the organisms. In obesity ILC2 numbers are reduced and their function is impaired, leading to the progression of metabolic inflammation. However, which events impact on ILC2 biology in the adipose tissue in obesity remains unresolved. Here, we find that high fat diet (HFD)-induced obesity in mice results in the metabolic reprogramming of adipose ILC2, impairing mitochondrial function and the expression of the enzyme Acetyl-CoA carboxylase 1 (ACC1). Investigating a possible connection between ACC1 and obesity-induced changes in ILC2, we show that fatty acids directly reduce the expression of ACC1, while pharmacological inhibition of ACC1 diminishes mitochondrial function and ILC2 metabolism. Furthermore, deletion of ACC1 in ILC2 phenocopies the overall reduction and functional impairment of ILC2 observed in obesity, which ultimately leads to increased triglycerides in circulation, adipose tissue hypertrophy and inflammation, even in the absence of HFD. Through single-cell RNA sequencing analysis we uncover that HFD-feeding or deletion of ACC1 results in the accumulation of undifferentiated ILC2 and ILC progenitors in the adipose tissue, suggesting that ACC1 may primarily regulate the maturation of ILC2. Together, these results reveal that obesity could predominately impair adipose ILC2 differentiation and activation by impacting on the expression of ACC1, rather than inducing cell death through lipid overload and lipotoxicity.

## Introduction

Rapid lifestyle changes, often related to reduced physical activity and increased consumption of energy-dense processed foods, have dramatically increased the risk of developing metabolic diseases, including obesity and diabetes, worldwide (Bluher, 2019). While various environmental and genetic factors contribute to the onset of obesity-related insulin resistance, the primary factor is often attributed to persistent low-grade inflammation in adipose tissue (Hotamisligil, 2006). Indeed, metabolic disorders coincide with fundamental changes in the adipose tissue (Hotamisligil, 2017). Healthy and non-obese adipose tissue-infiltrating immune cells consist mainly of adipose tissue macrophages (ATM), eosinophils, T regulatory cells (Tregs) and type 2 innate lymphoid cells (ILC2). This homeostasis is perturbed in obesity in mice and humans, where neutrophils, ILC1, NK cells, inflammatory macrophages and activated T cells comprise the majority of immune infiltrates. Furthermore, high-fat diet (HFD)- induced obesity in mice or obesity in humans correlates with a reduction of Tregs, ILC2 and ATM but an increase in metabolic inflammation (Brestoff et al., 2015; Cipolletta et al., 2012; Hams et al., 2013; Hildreth et al., 2021; Lee et al., 2015; Molofsky et al., 2013).ILC2 are a subset of innate lymphoid cells, a group of tissue resident immune cells found in mice and humans dedicated to the maintenance and defense of the body’s barrier sites, such as the skin, lung and intestine (Hazenberg and Spits, 2014; Spits et al., 2013; Vivier et al., 2018). Among the first important findings linking type 2 immunity to adipose tissue metabolism was the realization that mice lacking eosinophils exhibit increased weight gain (Wu et al., 2011). More recent data suggest that ILC2 expressing the transcription factor GATA-3 may maintain metabolic health of the organism through the recruitment of eosinophils and maintenance of ATM via the production of the cytokines IL-4, IL-5, and IL-13 (Hams et al., 2013; Molofsky et al., 2013; Nussbaum et al., 2013). Obesity results in the loss and functional impairment of adipose tissue ILC2, characterized by a substantial diminished capacity to produce IL-5 and IL-13 (Hams et al., 2013; Molofsky et al., 2013; Nussbaum et al., 2013; Oldenhove et al., 2018). Furthermore, experimental depletion of ILC2 or genetic ablation of IL-33, the cytokine pivotal for ILC2 activation, results in loss of eosinophils, ATM, weight gain, decreased glucose tolerance and insulin resistance, whereas treatment with IL-33 and IL-25, both activating ILC2, can reverse such effects and limit metabolic disease (Brestoff et al., 2015; Hams et al., 2013; Molofsky et al., 2013; Wilhelm et al., 2017). In addition, both co-stimulation of ILC2 through glucocorticoid-induced tumor necrosis factor receptor (GITR) (Galle-Treger et al., 2019) or stimulation with Death Receptor 3 (DR3) activating ILC2 to produce IL-13 (Shafiei-Jahani et al., 2020) ameliorates type 2 diabetes. Thus, the recent discovery of ILC2 as metabolically active cells and their dietary regulation offers a new approach to unravel their function in metabolic inflammation. However, despite the recent advances, it remains unresolved which events cause the loss of ILC2 in adipose tissue. A prerequisite for a better understanding of how dietary changes can lead to the disruption of the ILC2 compartment in adipose tissue is to uncover the metabolic constraints that fuel and maintain ILC2 function. Depending on the context, several metabolic pathways have been discovered to govern the regulation of ILC2 biology, including those associated with glucose, amino acid (AA), or fatty acid (FA) metabolism (Hodge et al., 2023; Monticelli et al., 2016; Panda et al., 2022; Surace et al., 2021; Wilhelm et al., 2016). Our previous work unraveled that upon activation, ILC2 strongly increase their capacity to acquire external lipids in allergen-driven airway inflammation (Karagiannis et al., 2020). Externally acquired lipids are converted into neutral triglycerides and stored in lipid droplets. This process is mediated by the gene catalyzing the conversion of diacylglycerol and fatty acyl CoA to triglycerides (TG) (*Dgat1*), serving the essential metabolic role of preventing lipotoxicity by funneling free fatty acids (FFA) within the cells into lipid droplets (LD). Both, expression of DGAT1 and increased external lipid acquisition are controlled by PPARγ and critical for the increase of ILC2 in chronic inflammation (Karagiannis et al., 2020). Moreover, PPARγ was shown to be important for IL-33-induced expansion of ILC2 within tumors (Ercolano et al., 2021) or the adipose tissue (Fali et al., 2021). Hence, expression of DGAT1, PPARγ and other genes important for lipid metabolism within cells may be particular important to allow tissue residency in lipid rich environments, such as the inflamed lung, tumor or adipose tissues. Following the general idea that understanding adipose tissue ILC2 metabolism may unravel cues explaining their dysfunction in metabolic diseases, in this study we found that ILC2 in obesity are metabolically impaired. Building upon the notion that understanding adipose tissue ILC2 metabolism may uncover explanations for their dysfunction in metabolic diseases, we found that ILC2s are metabolically impaired in obesity. Metabolic impairment is driven by increased lipid content present in obese mice and directly results in diminished expression of Acetyl-CoA carboxylase 1 (ACC1), an enzyme catalyzing fatty acid synthesis. Deletion of ACC1, but not DGAT1 or PPARγ in ILC2 leads to an overall reduction in adipose ILC2 numbers and function, which ultimately results in adipose tissue hypertrophy, increased circulating triglycerides, and inflammation of adipose tissue. Furthermore, single-cell RNA sequencing analysis, reveals that HFD or deletion of ACC1 prevents ILC2 differentiation, mediating the accumulation of ILC progenitors in the adipose tissue. Collectively, these results reveal that obesity, can impair type 2 immune cell differentiation and activation through ACC1- mediated metabolic reprogramming.

## Results

### ILC2 are metabolically impaired in obesity

Our previous work established that dietary cues regulate the function of ILC2 and control cellular metabolism (Karagiannis et al., 2020; Spencer et al., 2014; Wilhelm et al., 2016). To better understand the metabolic control of visceral adipose tissue (VAT) ILC2 at steady-state and in obesity, we placed mice on control or high fat diet (HFD) (60% kJ from fat), for a period of 12 weeks. As previously described, mice on a HFD displayed higher body and adipose tissue weight, as well as adipocyte hypertrophy, as assessed by hematoxylin and eosin (H&E) staining, confirming our diet-induced obesity model (**Figure 1A-C**). In addition, HFD feeding cause a decreased in adipose tissue ILC2 (CD45^+^Lin^-^Thy1.2^+^GATA3^+^), a markedly reduced expression of the cytokines IL-5 and IL-13 and a consequential reduction of eosinophils (**Figure 1D-F**). In addition, HFD reduced the percentages and absolute numbers of ATM marked by the expression of TIM4 (Cox et al., 2021), while inflammatory macrophages identified by expression of CD11c and absence of TIM4 infiltrated the adipose tissue (**Figure 1G-H**). Besides the functional alterations in ILC2 in the context of HFD-feeding, we also observed a shift in the metabolic profile of adipose ILC2 using a flow-cytometric based method for metabolic profiling of immune cells (Arguello et al., 2020). In this assay, inhibition of glucose usage and mitochondrial function through 2-deoxy-d-glucose (2-DG) and oligomycin, respectively is used to determine the usage and dependance on glucose and mitochondria (**Figure 1I**). Using protein synthesis measured by integration of puromycin as an overall assessment of metabolic activity, we found a decrease in puromycin uptake in ILC2 of HFD-fed mice compared to those in the control group (**Figure 1J**). We further observed a reduction in glucose dependance (tested through inhibition with 2-DG) but an increased capacity to metabolize FA and AA, while the mitochondrial dependence and glycolytic capacity (tested by inhibition with oligomycin) remained unchanged (**Figure 1K**). Moreover, both mitochondrial mass and potential, assessed by staining with Mitotracker green or tetramethylrhodamine methyl ester perchlorate (TMRM), respectively, were decreased (**Figure 1L-M**). This suggests that ILC2 in HFD-induced obesity may switch towards increased FA or AA metabolism, which as a consequence could result in mitochondrial impairment.

**Figure 1.**
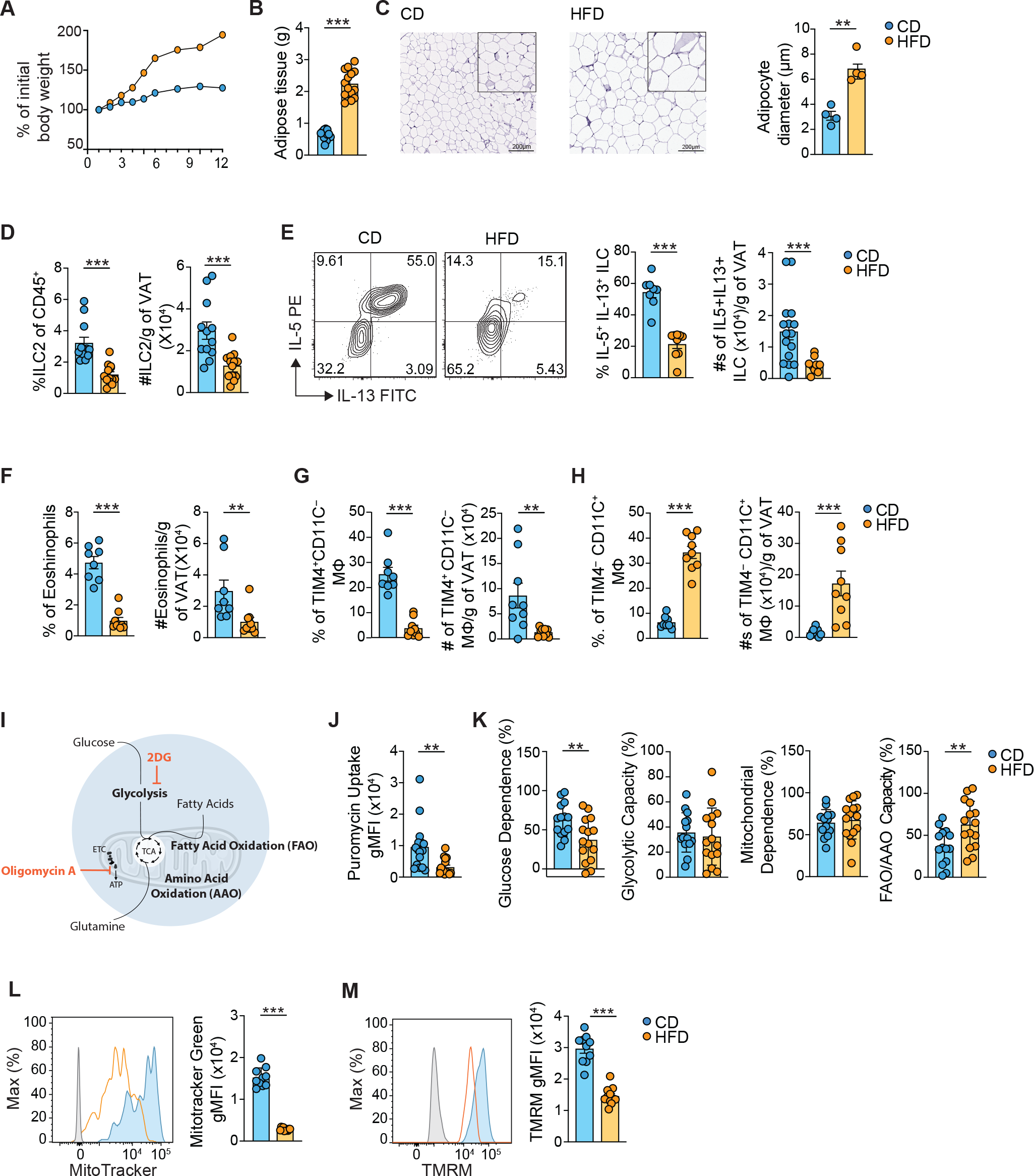
HFD-induced obesity impairs the metabolism of adipose ILC2 C57BL/6 mice were fed with either control (CD) or HFD diet for 12 weeks, euthanized and analysed. **(A)** Percentage of initial body weight and (**B)** weight of total VAT tissue. (**C)** Representative image of VAT tissues stained by hematoxylin and eosin (H&E) and quantification of the diameter of adipocytes. (**D)** Percentage and total numbers per gram of VAT of ILC2 (CD45^+^ Lin^−^ Thy1.2^+^ GATA3^+^). (**E)** Representative flow plots and percentage and total numbers per gram of VAT of IL-5^+^IL-13^+^ ILC (Lin^−^ Thy1.2^+^). (**F)** Percentage and total numbers per gram of VAT of eosinophils (CD45^+^MHC11^-^SiglecF^+^), (**G)** Percentage, total numbers per grams of VAT and total counts of TIM4^-^CD11C^+^ and (**H)** TIM4^+^CD11C^+^ macrophages. (**I)** Schematic of inhibitor function used for flow-based metabolic assessment. (**J)** gMFI values of puromycin uptake by VAT ILC2 and (K) analysis by flow-based metabolic assessment of VAT ILC2 (Lin^−^ Thy1.2^+^ GATA3^+^). (**L)** Representative histograms and gMFI values of MitoTracker green and (**M)** TMRM in ILC2. Results display one representative experiment **(A, C)** from at least two or two pooled experiments **(D-H, L, M)** or three **(J, K)** independent experiments with three to five mice in each experimental group. All graphs display means ± SEM; not significant (not shown) p>0.05 **p ≤ 0.01, *** p≤0.001. Scale bar represents 200μm

### Obesity controls ACC1 and fatty acid synthesis

To further understand the metabolic restrains imposed on adipose tissue ILC2 in the context of HFD that may lead to their reduction, we tested for the expression of key enzymes regulating essential metabolic processes. Reduced dependence on glucose was further corroborated by a reduction in the glucose transporter GLUT1 (**Figure 2A**). We did not detect any change in protein expression of carnitine palmitoyltransferase (CPT) 1a regulating fatty acid oxidation (FAO) (**Figure 2A**). However, enzymes involved with excess lipid metabolism, such as DGAT1 and PPARγ were decreased (**Figure 2A**). We also found a reduction in ACC1, the rate limiting enzyme encoded by *Acaca* and responsible for the first step in *de novo* fatty acid synthesis (FAS) by catalyzing the conversion of acetyl-CoA to malonyl-CoA (**Figure 2A**). Of note, malonyl-CoA inhibits CPT-1a and thus FAO (McGarry et al., 1977). Since ACC1 itself is inhibited by excess palmitate, ACC1 is an important switch point between FAO and FAS, and decreased expression in obesity could facilitate oxidation of excess lipids. Indeed, increased lipid exposure resulted in a downregulation of ACC1 in adipose ILC2 in vitro (**Figure 1I-K and Figure 2B**). To further test the role of FAO in ILC2 from obese mice, we adapted the initial metabolic flow-cytometric based assay to allow for the specific testing of FAO through addition of etomoxir, an inhibitor of FAO (**Figure 2C**). In line with an essential function of ACC1 regulating FAO, ILC2 from obese mice displayed increased dependance on FAO (**Figure 2D**). However, while the dependency and capacity to metabolize glutamine, assessed by addition of BPTES remained unchanged, their capacity to metabolize FA was reduced (**Figure 2D**). This suggests that ILC2 in obesity adapt towards FA as their major fuel, which may ultimately impact their mitochondrial capacity, explaining the observed decreased mitochondrial mass and potential (**Figure 1L, M**). Finally, we examined whether inhibition of ACC1 with 5- tetradecyloxy-2-furoic acid (TOFA) could cause metabolic disturbances, explaining the observed functional defects in ILC2. Indeed, pharmacological inhibition of ACC1 resulted in a drop of mitochondrial mass and potential, as well as a reduction in basal and the maximal rate of mitochondrial respiration, linking ACC1 function to mitochondrial activity (**Figure 2 E, F**).

**Figure 2.**
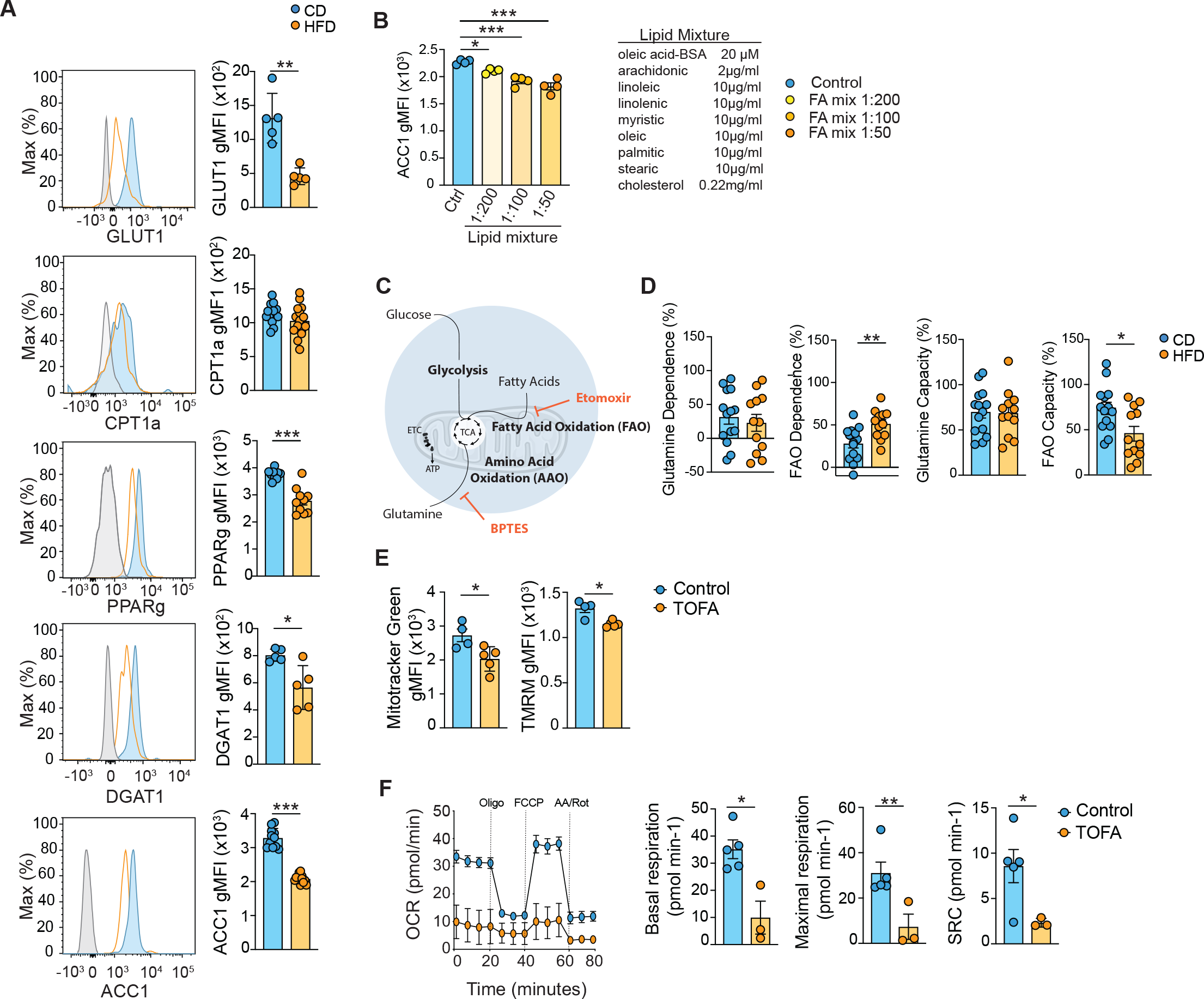
HFD-induced obesity reduces ACC1 and increases FAO in VAT ILC2. C57BL/6 mice were fed with either control or HFD diet for 12 weeks, euthanized and analysed. (**A)** Representative histograms and gMFI values of GLUT1, CPT1a, PPARγ, DGAT1 and ACC1 protein expression by VAT ILC2 (CD45^+^ Lin^−^ Thy1.2^+^ GATA3^+^). **(B)** gMFI values of ACC1 protein expression in VAT ILC2 cultured overnight without or with increasing concentrations of free fatty acids (FFA). Lipid mixture consists of 20 μM oleic acid-BSA together with a lipid cocktail containing 2μg/ml arachidonic and 10μg/ml each linoleic, linolenic, myristic, oleic, palmitic and stearic as well as 0.22mg/ml cholesterol. **(C)** Schematic of inhibitor function, **(D)** Analysis of puromycin integration in VAT ILC2 in dependance on glutamine (inhibition with BPTES) or fatty acid oxidation (inhibition with Etomoxir). **(E)** Energy metabolism analyzed by extracellular flux analysis on sort purified ILC2 in control and TOFA treatment 10uM. Oligo, oligomycin; AA/Rot, antimycin A and rotenone; OCR, oxygen consumption rate. (**F**) Representative histograms and gMFI values of MitoTracker green and TMRM in ILC2. Results display two pooled experiments **(A),** one representative experiment of at least two **(A, B, E, F)** or three pooled experiments independent experiments (**D**) with three to five mice in each experimental group. All graphs display means ± SEM; not significant (not shown) p>0.05, * p≤ 0.05, **p ≤ 0.01, *** p≤0.001.

### Deletion of ACC1 impairs the function of adipose ILC2, inducing adipose tissue inflammation

To assess a potential functional role of ACC1 in the metabolic regulation of adipose ILC2, we generated mice carrying a specific deletion of ACC1 in ILC2. This was achieved by crossing mice expressing the *RFP* gene followed by an internal ribosomal entry site (IRES) and the *CRE* recombinase under the endogenous control of the IL-5 locus (*Red5* mice, (Nussbaum et al., 2013)) to mice carrying the loxP-flanked *Acc1* gene (*ACC1^flox/flox^* mice). Since IL-5 is primarily produced by ILC2 in the steady state, such mice will specifically delete the target genes within the ILC2 compartment. Indeed, RFP expression was largely confined to ILC2 and 85-90% of RFP-positive cells in the adipose tissue were ILC2 identified by the expression of IL-33R (**Figure 3A**). Analysis of transgenic mice revealed that the frequency, total numbers and cytokine production of adipose tissue ILC2 from *Red5*/+ *x ACC1^flox/flox^*mice (termed ILC2^ACC1^) was substantially impaired in contrast to Red5/+ control mice, even in the context of normal chow feeding (**Figure 3B, C**). In contrast, when we generated mice carrying a specific deletion of Dgat1 and Pparg in ILC2, the frequency, total numbers and cytokine production of adipose tissue ILC2 in ILC2^ΔDgat1^ and ILC2^ΔPparg^ mice remained unchanged, excluding a general involvement of genes related to anabolic lipid metabolism in ILC2 (**Figure 3D, E**).

**Figure 3.**
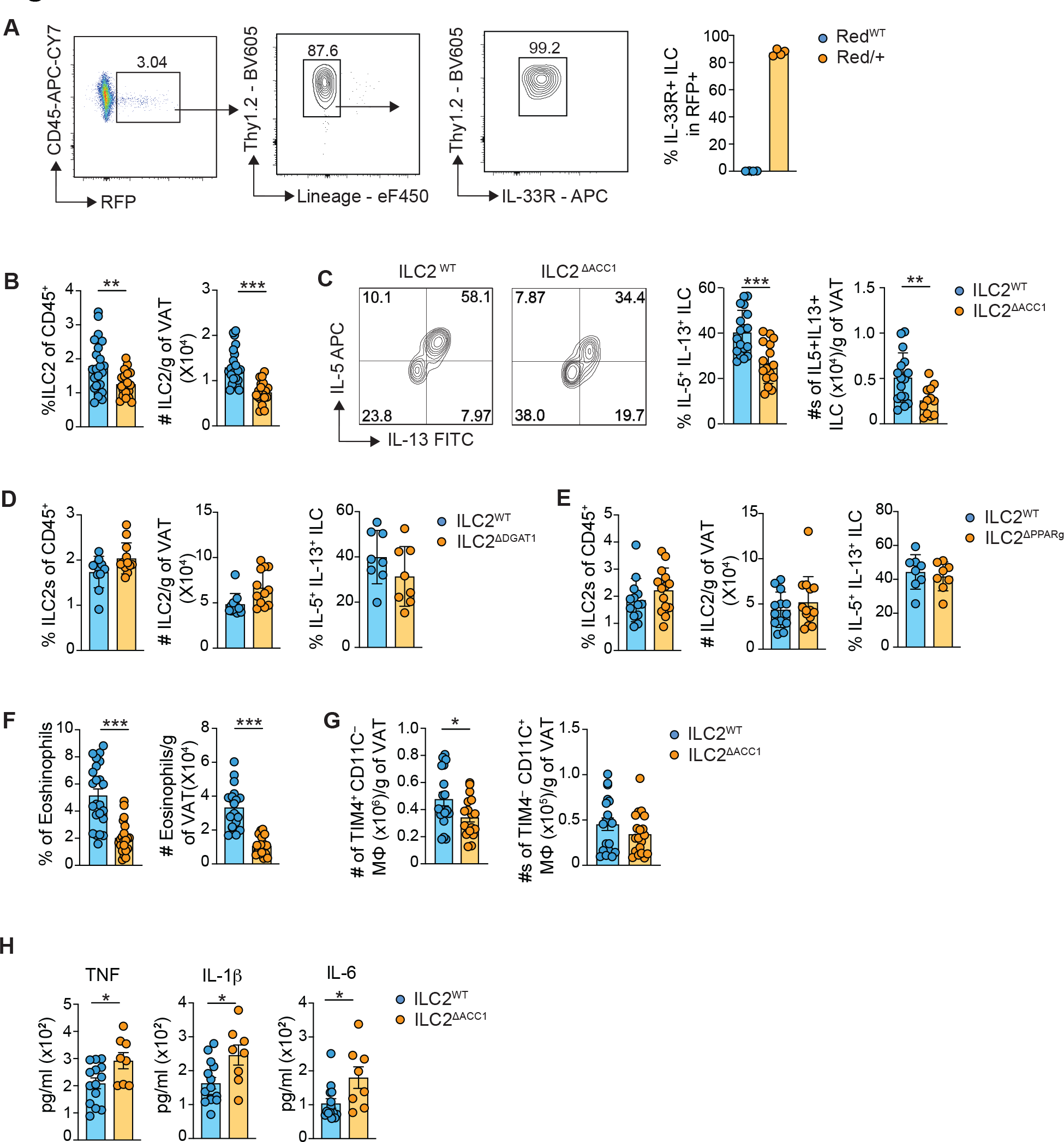
ACC1 deletion functionally impairs VAT ILC2 **(A)** Representative flow plots and percentage of ILC2 of RFP^+^ cells in Red/+. WT (ILC2) and ILC2^ΔACC1^ mice were euthanized and analyzed. **(B)** Percentage and total numbers per gram of VAT of ILC2, **(C)** Representative flow plots, percentage and total numbers of IL-5^+^IL-13^+^ per gram of VAT ILC. **(D)** Percentage and total numbers per gram of VAT of ILC2 and percentage of IL-5^+^IL-13^+^. **(E)** Percentage and total numbers per gram of VAT of ILC2 and percentage of IL-5^+^IL-13^+^ **(F)** Percentage and total numbers per gram of VAT of eosinophils MHCII^−^ Siglec F^+^. **(G)** Total numbers per gram of VAT of TIM4^+^CD11C^-^ and TIM4^-^CD11C^+^ macrophages. (**H)** Concentration of cytokines in the VAT homogenate. Results display one representative experiment of at least two **(A, H)**, two or three pooled experiments **(B-G)** with three to five mice in each experimental group. All graphs display means ± SEM; not significant (not shown) p>0.05, * p≤ 0.05, **p ≤ 0.01, *** p≤0.001.

ILC2 are essential for maintaining metabolic health and adipose tissue homeostasis via the promotion of type 2 immunity and the recruitment of eosinophils and activation of ATM (Hams et al., 2013; Molofsky et al., 2013; Nussbaum et al., 2013). Hence, we hypothesized that impaired metabolic maintenance of ILC2 in the absence of ACC1 might result in defective type 2 immunity leading to enhanced adipose tissue inflammation. Indeed, we observed a significant reduction of both, adipose tissue eosinophils and TIM4+ tissue resident macrophages (**Figure 3F, G**). In contrast, the numbers of pro-inflammatory macrophages remained unchanged in ILC2^ΔACC1^ mice in comparison to ILC2^WT^ control mice (**Figure 3G**). Next, we examined the production of inflammatory cytokines. Consistent with impaired type 2 immunity, levels of the inflammatory cytokines TNF, IL-1β and IL-6, which have previously been shown to drive adipose tissue inflammation (de Baat et al., 2023; Hotamisligil, 2017), were significantly increased in the adipose tissue of ILC2^ΔACC1^ mice (**Figure 3H**). Thus, expression of ACC1 is essential for the maintenance and function of VAT ILC2 to prevent perturbance of adipose tissue immune homeostasis and inflammation.

### Deletion of ACC1 in ILC2 alters adipose metabolism

In light of the profound changes in the composition of immune cells in VAT and the increased expression of inflammatory cytokines, we next examined a potential effect of disrupted ILC2 function on adipose tissue metabolism in ILC2^ΔACC1^ mice. Notably, deletion of ACC1 in ILC2 and a consequential loss of VAT ILC2 resulted in increased adipose tissue weight, leading to adipocyte hypertrophy assessed by histological analysis (**Figure 4A, B**). Furthermore, ILC2^ΔACC1^ mice displayed increased triglycerides in circulation (**Figure 4C**), indicating a shift towards heightened lipogenesis. Indeed, genes involved in adipose tissue lipogenesis, including *Plin2*, *Pparg, Dgat1*, and *Adipoq* or cholesterol synthesis such as *Srebf1* were significantly increased in the VAT of ILC2^ΔACC1^ mice (**Figure 4D**). In addition to the morphological changes and the increased adipose tissue inflammation, we also observed increased expression of fibrotic genes *Col3a*, *Acta2* and *Ctgf* **(Figure 4E**). Taken together, these data indicate that loss of ACC1 in ILC2 induces a shift towards lipogenesis and fibrotic remodeling of the adipose tissue.

**Figure 4.**
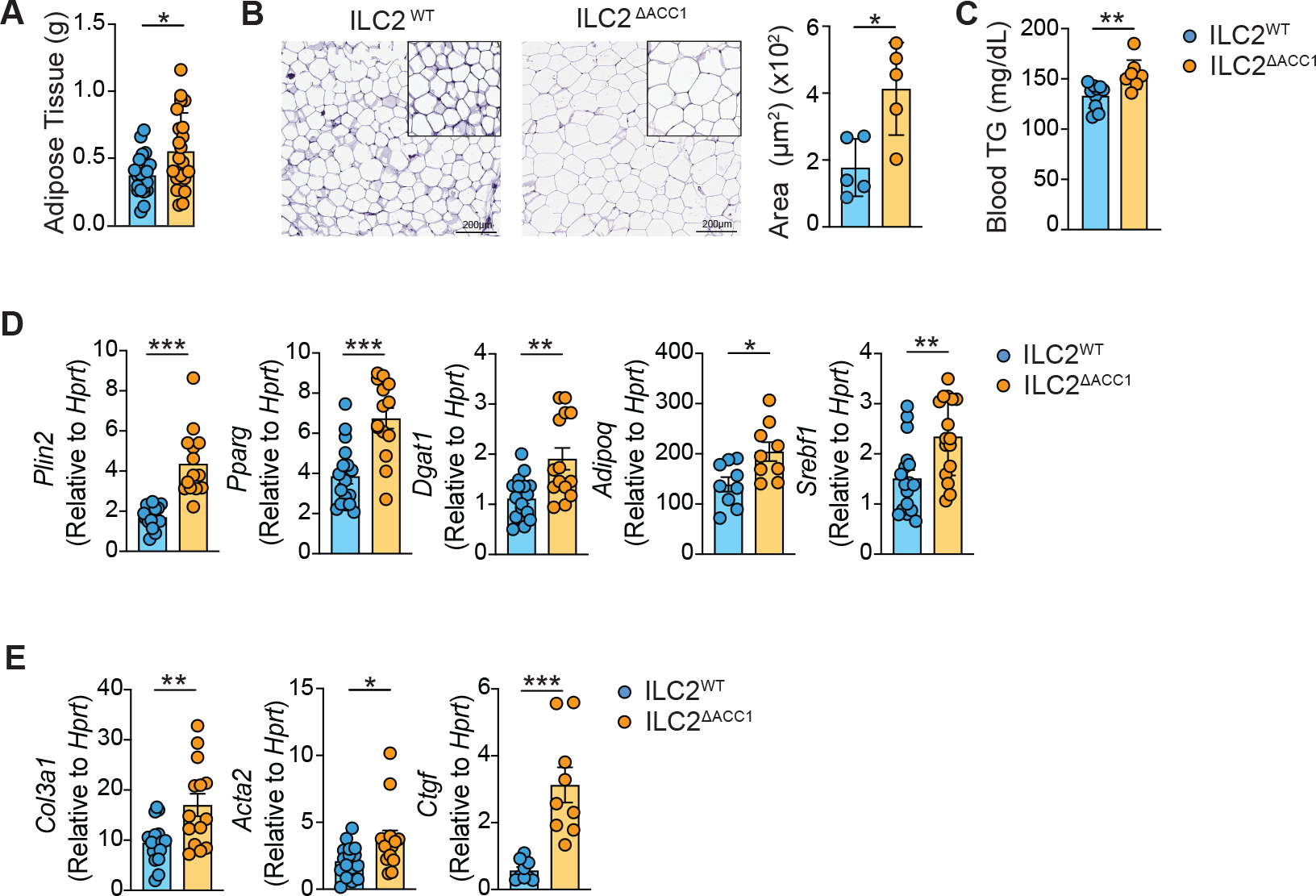
ACC1 deletion induces adipose tissue inflammation and fibrosis. WT (ILC2) and ILC2^ΔACC1^ mice were euthanized and analyzed. **(A)** Weight of total VAT tissue. (**B)** Representative image of VAT tissues stained by hematoxylin and eosin (H&E) and quantification of the area of adipocytes. (**C)** Concentration of triglycerides in serum. **(D-E)** RT-PCR analysis relative to *Hprt* of VAT tissue. Results display three pooled independent experiments **(A, D, E)** or one representative experiment of at least three **(B, C)** independent experiments with three to five mice in each experimental group. All graphs display means ± SEM; not significant (not shown) p>0.05, * p≤ 0.05, **p ≤ 0.01, *** p≤0.001. Scale bar represents 200μm.

### Obesity prevents the differentiation of adipose ILCp into ILC2

To further understand the control of ILC2 in obesity, we analyzed a publicly available single cell sequencing dataset of adipose tissue lymphocytes from lean and HFD-fed mice (Yang et al., 2022). Due to the highly similarity between ILC2 and T, NK cells, we first extracted cells labeled as T, NK and ILC from the dataset to determine ILC2 related populations. Cluster analysis of these extracted cells enabled us to find ILC2 progenitors (ILCp) based on high expression of genes *Zbtb16*, *Tcf7* and *Il18r1* as well as intermediate and mature ILC2 characterised by the expression of *Il1rl1*, *Gata3*, *Il13, Il5* and *Klrg1* (**Figure 5A, B**). In line with our *in vivo* data, quantification of mature ILC2 in lean and HFD condition indicated that HFD-feeding reduced numbers of mature ILC2 and of ILC2 in an intermediate cellular state (**Figure 5A, B**). To explore the impact of HFD on ILC2 differentiation, we conducted trajectory analysis to infer ILC2 development process. This analysis unveiled a potential hindrance in the differentiation potential of adipose ILC2 (**Figure 5C**). In fact, alongside the general reduction in cytokine-producing ILC2, we observed an accumulation of a recently described cell type termed tissue-resident ILC progenitors (ILCp) in the lung (Zeis et al., 2020) (**Figure 5D**). Tissue ILCp are marked by the absence of lineage markers, GATA-3 and IL-33 receptor (IL-33R) but express IL-18Rα. Indeed, by staining for the absence of lineage markers and GATA-3 but the expression of IL-18Rα, we confirmed accumulation of ILCp in the adipose tissue of HFD mice, while the number of cytokine-producing mature ILC2 was reduced (**Figure 5D, E**).

**Figure 5.**
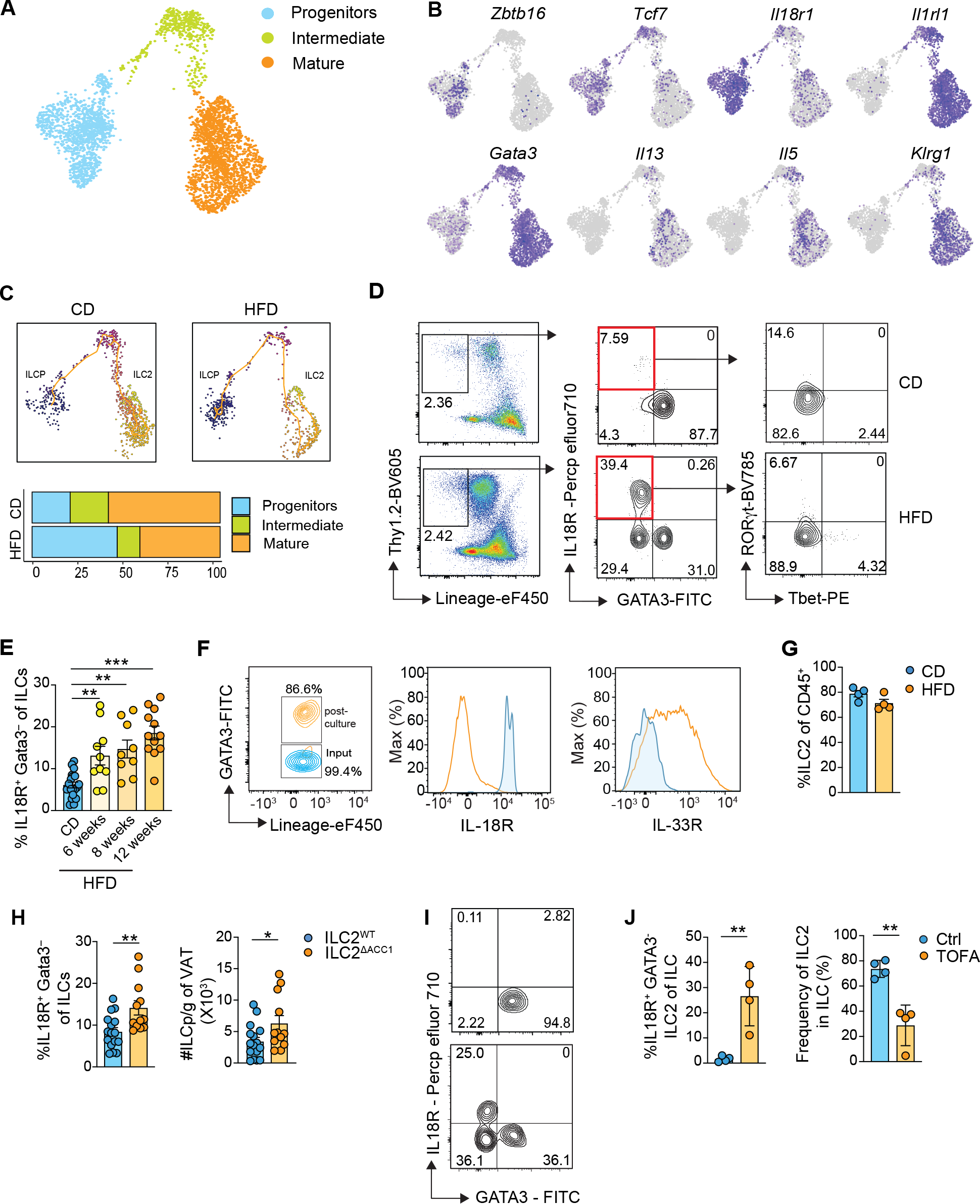
Obesity prevents the differentiation of VAT ILCp to ILC2 by impairing expression of ACC1. Single cell transcriptomic analysis of adipose tissue ILC2 from lean and HFD-fed mice. **(A)** UMAP embedding of ILC2 clusters. Data were extracted from published single cell RNA-seq dataset. **(B)** Feature plot of canonical genes expressed in ILC2 progenitors and mature ILC2 cells. **(C)**Trajectory analysis of ILC progenitor differentiation to ILC2 and bar chart of population frequency in CD and HFD. Representative flow plots **(D)** and percentage **(E)** of IL18R^+^GATA3^−^ ILCP from adipose tissues of mice fed with control diet for 12 weeks or HFD diet for 6, 8 or 12 weeks. (**F)** Representative flow plots of sort purified adipose ILCP (CD45^+^ Lin^−^ Thy1.2^+^ IL18R^+^IL33R^−^) before (“input”) and after 14 days culture (“output”) in the presence of OP9 cells and IL7, IL25, IL33 and mSCF cytokine mix. (**G)** Percentage of ILC2 (CD45^+^ Lin^−^GATA3^+^) of total cultured cells. **(H)** Percentage and total numbers per gram of VAT of ILCP from WT (ILC2) and ILC2^ΔACC1^ mice. **(I)** Representative flow plots of bone marrow CLP (CD45^+^Lin^−^a_4_b_7_^−^IL-7Ra^+^CD90.2^−^) after 14-days co-culture with OP9 cells and cytokine mix IL7, IL25, IL33 and mSCF in the presence of absence of TOFA. **(J)** Percentage of IL18R^+^GATA3^-^ ILCP and ILC2 (CD45^+^ Lin^−^GATA3^+^) in total ILC after bone marrow CLP differentiation in the presence or absence of TOFA. Results display two pooled experiments **(E)**, one representative experiment of two **(F, G, I, J)** or three pooled **(H)** independent experiments with three to five mice in each experimental group. All graphs display means ± SEM; not significant (not shown) p>0.05, * p≤ 0.05, **p ≤ 0.01, *** p≤0.001.

Lung resident ILCp were shown to be able to differentiate into mature ILC2. In order to confirm that adipose tissue resident ILCp possessed similar potential, we co-cultured adipose ILCp with OP-9 cells expressing the Notch ligand delta. Consistent with previous results in the lung, culture of ILCp together with IL-7, IL25, IL-33 and stem cell factor (SCF) resulted in the differentiation of mature GATA3^+^ ILC2, further confirming the progenitor potential of adipose resident ILCp (**Figure 5F**). However, we did not observe a difference between the potential of ILCp isolated from Ctrl or HFD mice to differentiate into mature GATA-3 expressing ILC2, further confirming an impact of HFD on ILC maturation but not progenitor potential (**Figure 5G**).

Given the substantial reduction in ILC2 and potential to produce cytokines in ILC2^ΔACC1^ mice, we next examined the importance of ILC2-intrinsic ACC1 for tissue specific differentiation and maturation. Consistently, with a role of ACC1 in the differentiation we observed increased percentages and numbers of ILCp in ILC2^ΔACC1^ mice (**Figure 5H**), Finally, we confirmed a direct effect of ACC1 in the differentiation also of bone marrow (BM) ILCp into ILC2 by pharmacological blockade of ACC1 with TOFA. While BM common lymphocyte progenitors (CLP) differentiated almost exclusively into mature ILC2 in the presence of IL-7, IL-25, IL-33 and SCF, blocking ACC1 had a profound effect on ILC2 differentiation and resulted in a significant increase in the percentage of IL-18Rα immature tissue ILCp at the expense of GATA-3^+^ mature ILC2 (**Figure 5I, J**). Collectively, these results strongly imply that HFD and impaired expression of ACC1 impact on the differentiation of ILCp into mature ILC2 in the adipose tissue.

## Discussion

Analyzing the metabolic constrains governing the control of adipose tissue ILC2, we find that ACC1 represents an integral component of the bioenergetic pathways controlling the maintenance of adipose tissue ILC2. In obesity, the metabolic balance of ILC2 is perturbed resulting in a switch towards increased FAO to maintain their bioenergetic demand. However, this metabolic alteration, likely prompted by increased lipid availability, may in the long run result in a decline in mitochondrial metabolic activity, as evidenced by a significant decrease in both mitochondrial mass and potential. Consequently, the development of ILC2 is hampered, resulting in a pronounced reduction in mature, cytokine-producing ILC2, while ILC progenitors accumulate in the adipose tissue. Our findings suggest that this metabolic shift in ILC2 cells may be orchestrated by the downregulation of ACC1. Deleting ACC1 specifically in ILC2 cells phenocopies several aspects observed in obesity, including impaired recruitment of eosinophils and reduced maintenance of TIM4^+^ ATM. This deficiency in ILC2-mediated type 2 immunity leads to heightened inflammation in adipose tissue, which consequently affects metabolic health, as evidenced by elevated levels of circulating triglycerides and exacerbated adipocyte ballooning. Our findings are significant in multiple aspects. We uncover that obesity, rather than solely increasing lipotoxicity, hampers the development of ILC2 cells by influencing ACC1 expression, offering a novel hypothesis regarding the regulation of cellular compartmental shifts. In this context, ACC1 emerges as a key player, modulating the balance between anabolic and catabolic lipid metabolism. Various roles of ACC1 in immune cells have been documented. For instance, ACC1 is implicated in controlling the development of pathogenic Th17 cells but not T regulatory cells (Tregs) (Berod et al., 2014). Notably, the reliance of Th17 cells on ACC1 expression seems to be crucial in obesity as well, since ACC1 is upregulated in Th17 cells in HFD-fed mice (Endo et al., 2015). Furthermore, inhibiting ACC1 also appears to impede the differentiation of DC (Nguyen-Phuong et al., 2022). Thus, the function of ACC1 could therefore be regulated differently in immune cells, depending on whether they have a positive or negative effect on metabolic health. Moreover, our results validate previous research emphasizing the crucial role of ILC2 in metabolic health, while also expanding on these findings by demonstrating that metabolic dysregulation, specifically in ILC2 cells induces alterations in adipose tissue immune cell composition, inflammation, and lipid metabolism. Previous results largely build on investigating the function of VAT ILC2 by exogenous delivery of IL-33 or deletion of IL-33 and IL-33R. Here, IL-33 expressed by stomal cells, including those found in the adipose tissue (Dahlgren et al., 2019; Mahlakoiv et al., 2019; Rana et al., 2019; Spallanzani et al., 2019) acts as a key regulator of adipose tissue immune homeostasis by maintaining ILC2, but also Tregs (Kolodin et al., 2015; Vasanthakumar et al., 2015). Hence, the investigation of the function of ILC2 in metabolic health in the context of IL- 33 biology was previously hampered by potential overlapping effects on Treg biology. By specifically targeting ACC1 in ILC2, we overcome some of the previously experienced potential limitation and further confirm the unique importance of ILC2 for the maintenance of adipose metabolism and metabolic health. Finally, connecting the underlying causes of metaflammation to potential alterations in immune cell metabolism is crucial, as certain adverse effects observed in treatment of diabetes might also affect immune cell metabolism. Thus, restoring adipose tissue metabolism may fail to address immune-mediated components of obesity and adopting a more comprehensive approach that considers immune cell metabolism could inform the development of effective treatments.

## Acknowledgements

We thank Dr. Richard Locksley for providing *Red5* mice and *Il1r1*^-/-^ mice; the animal facility staff for animal care at the UKB (HET); staff from the UKB flow core facility, Andreas Dolf, Peter Wurst, Praveen Mathoor, and David Kühne; the UKB sequencing core facility; the UKB bioinformatics facility; the UKB imaging facility and Alice Jacob and Prof. Dr. Zeinab Abdullah for sharing adipose tissue samples from dietary mouse studies. We thank the Wilhelm lab for critical discussions regarding the manuscript.

## Funding

This work was supported by the Ministry for Science and Education of North-Rhine-Westphalia, the Human Frontiers Science Program (HFSP) and the Deutsche Forschungsgemeinschaft (DFG, German Research Foundation) under Germany’s Excellence Strategy – EXC2151 – 390873048, the DFG collaborative research center – SFB 1454– Project-ID 432325352, and the DFG program grants (WI 4554/1-1 and WI 4554/6-1).

## Author contributions

M.R.T. designed and performed the experiments. J.Y. performed single cell sequencing analysis. F.N., C.W., P.W. and R.R. provided experimental help. J.Z. and C.T. performed the lipidomic analysis. F.K., J.H., and K.H. provided expertise in data analysis. C.W. designed the experiments and wrote the manuscript.

## Declaration of Interests

C.W. is a consultant for Odyssey Therapeutics and Orphagen Pharmaceuticals.

## Materials and Methods

### Animals

C57Bl/6 (WT) were bred in-house or purchased from Charles River labs and maintained at in-house facilities. *Red5* animals (kindly provided by Dr. Richard Locksley, University of California, San Francisco) were crossed to *ACC1^fl/fl^*, *Dgat1^fl/fl^* and *Pparg ^fl/fl^* (purchased from Jackson Laboratory) and maintained at in-house facilities. All mice were kept under specified pathogen free (SPF) conditions. All procedures were performed according to ethical protocols approved by the local and regional ethics committees. All mice used were between 6-10 weeks of age.

### Diet studies

Control diet D12450B LS (13kJ% Fat, low sugar) and high fat diet E15742-347 (60kJ% Fat) were purchased from Ssniff. Mice were placed on high fat or control diets at 6 weeks after birth and maintained on the appropriate diet for at least 12 weeks before sacrificing. In some experiments mice were sacrificed at earlier timepoints as indicated in the results.

### Cell isolation from visceral adipose tissue and flow cytometry

Visceral white adipose tissues were diced and digested with 0.2 mg/ml Liberase TL (Roche) and 0.5 mg/ml DNase I (Sigma) at 37°C for 25 minutes. For cell sorting and whole adipose culture experiments, isolated adipose tissue cells were further purified using a 37.5% Percoll gradient, followed by lysis of red blood cells with Ammonium-Chloride-Potassium (ACK) lysing buffer. Single-cell suspensions were stained with fluorochrome-conjugated antibodies against any combination of the following surface antigens: CD3, CD11b, CD11c, CD19, Ter119, NK1.1, Gr-1, CD45, Thy1.2, IL33R, IL18R, MHCII, Siglec F, F4/80, TIM4, a4b7 and IL7Ra.

DAPI or fixable Zombie UV dye (Biolegend) were used to exclude dead cells. For examination of transcription factors and cellular proliferation, cells were subsequently treated with the Foxp3 fixation/permeabilization kit (eBioscience) in accordance with the manufacturer’s instructions and stained for 60 min at 4°C with fluorochrome-conjugated antibodies against GATA3, RORγt, Tbet and Ki67. To assess the expression of metabolic enzymes, cells were stained intracellularly with unconjugated polyclonal antibodies against GLUT1, CPT1a, DGAT1, PPARγ and ACC1 for 60 min at 4°C, followed by 20 min staining with fluorochrome-conjugated goat anti-rabbit IgG secondary antibodies.

### Re-stimulation of cells for intracellular cytokine staining

Cells isolated from adipose tissue were stimulated for 3 h with Phorbol 12-myristate 13-acetate (0.5 µg/ml) (PeproTech) and ionomycin (0.5 µg/ml) (Sigma - Aldrich) in the presence of brefeldin A (1μg/ml) (GolgiPlug, BD Biosciences). Stimulated cells were stained for surface markers, fixed with 3.7% formaldehyde (Sigma), and stained with antibodies against IL-5, and IL-13 for 60 min at 4°C.

### Mitochondrial mass, membrane potential and uptake of FA by FACS

Mitochondrial mass, membrane potential, or FA uptake of freshly isolated or cultured adipose tissue cells were assessed by staining with 20nM TMRM (Sigma – Aldrich), 40nM MitoTracker Green (Thermo Fisher) and 25 ng/ml BODIPY FL C_16_ (ThermoFisher) respectively, for 30 min at 37°C in FBS free complete medium. Cells were washed twice, stained with surface antibodies, and analyzed by flow cytometry.

### Culture of adipose tissue lymphocytes

Freshly isolated lymphocytes from adipose tissue were cultured in 50 μl RPMI 1640 supplemented with 3% FBS, penicillin, streptomycin, HEPES, glutamine, nonessential amino acids, 50mM β-mercaptoethanol (Complete Media), and IL-7 (20 ng/ml, Biolegend) for 2 days. To test the effect of fatty acids on ACC1 expression, cells were cultured with a mixture of 20 μM oleic acid-BSA (Sigma) together with a lipid cocktail containing 2μg/ml arachidonic and 10μg/ml each linoleic, linolenic, myristic, oleic, palmitic and stearic as well as 0.22mg/ml cholesterol (Sigma) in final concentration as indicated in the results. To test the effect of ACC1 inhibition on ILC2 mitochondrial mass and membrane potential, freshly isolated adipose tissue cells were cultured with 10μM, TOFA (Cayman Chemicals). After 2 days of treatment cells were stained and analyzed by flow cytometry.

Fluorescence-activated cell sorting of ILC2, adipose tissue ILCP and bone marrow ILCp ILC2 were sorted by flow cytometry from total adipose tissue lymphocytes isolated from naïve or IL-33-challenged mice based on the absence of lineage markers (CD3, CD11b, CD11c, CD49b, CD19, Ter119, NK1.1, and Gr-1) but expression of Thy1.2, CD45, and IL33R. Adipose tissue ILCP were sorted from CD or HFD-fed mice based on the absence of lineage markers but expression of Thy1.2, CD45, and IL18R. BM ILCp isolated from naïve mice were identified and sorted as Lin– a4b7–IL-7Ra+CD90.2–.

### Preparation of OP9-DL4 feeder cell layer and cell culture of sorted ILCp

Feeder cells were grown in MEM Alpha medium (Gibco) supplemented with 10% FBS and 1% penicillin-streptomycin. Feeder cells were grown until 80% - 90% confluency and detached with trypsin-EDTA at 37°C for 3 min. Stromal cells were plated at a density of 10,000 cells per well in a flat bottom 96-well plate, grown overnight, followed by treatment with Mitomycin- C at 50μg/ml final concentration for 25 minutes. Cells were washed three times with PBS. Freshly sorted adipose ILCP or BM CLP (1000 cells/ well) were plated onto Mitomycin-C treated feeder cells and cultured in RPMI 1640 medium supplemented with 10% FBS, 1% penicillin-streptomycin and a cytokine mixture of IL-7 (20ng/ml), murine stem cell factor (SCF) (10ng/ml), IL-33 (10ng/ml) and IL-25 (10ng/ml). To assess the effects of ACC1 inhibition, BM CLP were cultured in the presence 10μM TOFA (Cayman Chemicals). Half of medium was removed and replaced by fresh medium every 3-4 days. Cells were analyzed by flow cytometry after 14 days.

### Administration of IL-33

WT C57Bl/6 mice were injected intraperitoneally with 250 ng recombinant mouse IL-33 (Biolegend) in 100 µl PBS for three consecutive days. Mice were euthanized 7 days after the first challenge and cells from the adipose tissue were isolated for analysis.

### Lipidomic analysis of ILC2

ILC2 isolated from adipose tissues were resuspended in 500 µl of extraction mix (490 μl MeOH/CHCl_3_ 5:1 and 10 μl internal standard mix containing alkyne-labeled lipids) and processed for extraction, as previously described by Thiele et al. (Thiele et al., 2019). The dissolved lipids were analyzed on a Thermo Q Exactive Plus spectrometer equipped with a standard heated electrospray ionization ion source using direct injection from a Hamilton syringe driven by a syringe pump under the control of the Tune instrument control software. Raw files were converted to .mzml files using MSconvert and analyzed using LipidXplorer for lipid species that incorporated the alkyne fatty acid

### Metabolic Assays

For real-time measurement of the oxygen consumption rate (OCR), purified ILC2 isolated from IL-33-challenged mice were cultured overnight with 10μM TOFA in complete media supplemented with 20 ng/ml IL-7 (Biolegend). Cells were washed with assay medium supplemented with 10 mM glucose, 1 mM pyruvate, and 2 mM glutamine and analyzed with an XF-96 Extracellular Flux Analyzer (Agilent). Four consecutive measurements were obtained under basal conditions followed by the addition of 1 μM oligomycin, which inhibits the mitochondrial ATP synthase; 1.5 μM FCCP (Carbonyl cyanide 4-(trifluoromethoxy) phenylhydrazone), which uncouples ATP synthesis from oxygen consumption, and a combination of 100 nM rotenone plus 0.5 μM antimycin A, which inhibit the electron transport chain by blocking complex I and III, respectively. All chemicals used for these assays were obtained from Sigma-Aldrich.

### Metabolic profiling of immune cells by flow

Mouse adipose tissue cells were incubated for 30 min at 37 °C, 5% CO_2_ followed by treatment for 15 min at 37 °C, 5% CO_2_ with control, 2-DG (100 mM; Sigma-Aldrich), oligomycin (1 μM; Sigma-Aldrich) or a combination of both drugs. Puromycin (10 μg ml^−1^; Abcam) was added for 60 min at 37 °C. After staining with primary antibodies as described above cells were fixed and permeabilized using Foxp3 fixation/permeabilization kit (eBioscience) following the manufacturer’s instructions. Intracellular staining of puromycin was performed by incubation with the anti-puro monoclonal antibody (1:1000, Clone MABE343, Merck) for 60 min at 4 °C. To assess the mitochondrial fuel usage, isolated adipose tissue lymphocytes were incubated for 30 min at 37 °C, 5% CO_2_ followed by treatment for 15 min at 37 °C, 5% CO_2_ with control, BPTES (3 μM; Cayman Chemical), Εtomoxir (4 μM; Cayman Chemical), 2-DG (100 mM; Sigma-Aldrich) or a combination of drugs. BPTES, Etomoxir and 2-DG to inhibit glutamine oxidation, long chain fatty acid oxidation and glucose oxidation, respectively. Puromycin (10 μg ml^−1^; Abcam) was added for 60 min at 37 °C. After staining with primary antibodies as described above cells were fixed and permeabilized using Foxp3 fixation/permeabilization kit (eBioscience) following the manufacturer’s instructions. Intracellular staining of puromycin was performed by incubation with the anti-puro monoclonal antibody (1:1000, Clone MABE343, Merck) for 60 min at 4 °C. The following equations were used to calculate the percentages of fuel dependence and capacity:

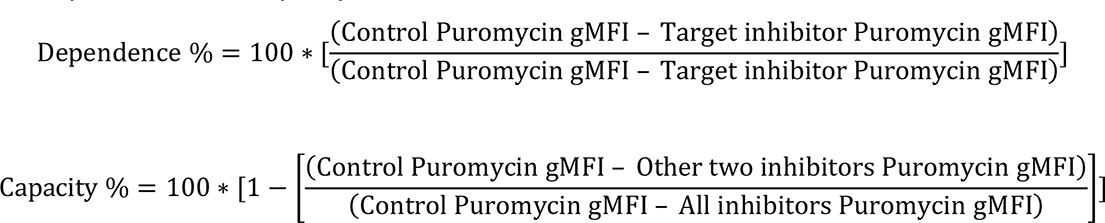

### Multiplex quantification of cytokines from visceral adipose tissue

For tissue cytokines detection, 5 mg of mouse VAT tissue was homogenized in 300 μl of RIPA lysis buffer (R0278-50ML, Sigma-Aldrich) containing protease inhibitor and centrifuged at 12,000g for 10min at 4 °C. The supernatant was collected and cytokine concentrations were measured by Legendplex (Biolegend) according to the manufacturer’s instructions.

### Histology and blood triglycerides

A piece of the visceral adipose tissue was collected and fixed in 10% formaldehyde solution and embedded into paraffin blocks. Sections were stained with hematoxylin and eosin (H&E). Quantification of adipocyte diameter was accomplished with Image J software and the open source Adiposoft plugin (Galarraga et al., 2012). Plasma triglycerides were measured with Cobas p 312 (Roche).

### Real-time PCR

RNA was extracted from purified VAT ILC2 or tissue samples using Trizol (ThermoFisher Scientific) and reverse transcribed with RevertAid Kit (Thermo Fischer Scientific) according to the manufacturer’s instructions. The cDNA served as a template for amplification of the genes of interest. For analysis of genes expressed by ILC2 and tissues, TaqMan probes for *Plin2, Pparg Dgat1, Adipoq, Srebf1, Col3a1, Acta2 and Ctgf* (IDT) were used and target-gene expression was calculated using the comparative method for relative quantification upon normalization to *Hprt* gene expression.

### Determination of ILC from publicly available dataset

Public scRNA dataset GSE183288 is downloaded from GEO dataset (Yang et al., 2022) and is immediately loaded into R (v4.1.3) To determine ILC progenitor and mature ILC2 populations, we first extracted cells labeled as T, NK and ILC from visceral white adipose tissue and then performed cluster analysis using Seurat (v4.1.0; (Hao et al., 2021). In brief, after normalization, scaling and PCA analysis on the downloaded dataset, the first 30 PCs were used to construct a shared nearest-neighbor graph. Cluster analysis with resolution of 0.5 was performed to generate 14 cell clusters. ILC progenitor cells and ILC2 were extracted and analyzed in Seurat with the same steps. After cluster analysis, the UMAP based dimension reduction was performed to visualize cells in low dimensions.

### Trajectory inference of ILC development

To infer the ILC development process, ILC cells determined from the last step were submitted for trajectory analysis. In short, the seurat object of ILC cells was first converted to cell_data_set object using SeuratWrappers (v0.3.2). After cluster analysis, a principal graph was generated to determine the location of development branches, and to estimate development pseudotime using Monocle 3 (v1.3.4;(Cao et al., 2019)).

### Data and code availability

R scripts used for the analysis have been deposited on GitHub (https://github.com/JiangyanYu/ILC2_Theodorou_2024). Scripts were executed in the docker image: https://hub.docker.com/r/jiangyanyu/jyu_r4.1.2.

### Quantification and Statistical Analysis

Data were analyzed with the Prism software (GraphPad). A two-tailed Student’s t test was used for all statistical analyses: * p≤ 0.05, **p ≤ 0.01, *** p≤0.001. Statistical details are indicated in the figure legends.

